# *Sox21b* underlies the rapid diversification of a novel male genital structure between *Drosophila* species

**DOI:** 10.1101/2023.08.17.552955

**Authors:** Amber M. Ridgway, Emily Hood, Javier Figueras Jimenez, Maria D. S. Nunes, Alistair P. McGregor

## Abstract

The emergence and subsequent diversification of morphological novelties is a major feature of animal evolution^1–9^. However, in most cases little is known about the molecular basis of the evolution of novel structures and the genetic mechanisms underlying their diversification. The epandrial posterior lobes of the male genital arch is a novelty of some species of the *Drosophila melanogaster* subgroup^10–13^. The posterior lobes grasp the ovipositor of the female and then integrate between her abdominal tergites, and therefore these structures are important for copulation and species-recognition^10–12,14–17^. The posterior lobes evolved from co-option of a Hox regulated gene network from the posterior spiracles^10^ and have since diversified in shape and size in the *D. simulans* clade in particular over the last 240,000 years driven by sexual selection^18–21^. The genetic basis of this diversification is highly polygenic but to the best of our knowledge none of the causative genes have yet been identified despite extensive mapping^22–30^. Identifying the genes underlying the diversification of these secondary sexual structures is essential to understanding the basis of changes in their morphology and the evolutionary impact on copulation and species recognition. Here, we show that the transcription factor encoded by *Sox21b* negatively regulates posterior lobe size during development. This is consistent with higher and expanded expression of *Sox21b* in *D. mauritiana*, which develops smaller posterior lobes compared to *D. simulans*. We tested this by generating reciprocal hemizygotes and confirmed that changes in *Sox21b* underlie posterior lobe evolution between these two species. Furthermore, we found that differences in posterior lobe size caused by the species-specific allele of *Sox21b* significantly affect the duration of copulation. Taken together, our study reveals the genetic basis for the sexual selection driven diversification of a novel morphological structure and its functional impact on copulatory behaviour.

**Highlights:**

- *Sox21b* regulates the development of the epandrial posterior lobes, a recently evolved novel structure of some species of the *Drosophila melanogaster* subgroup, which has subsequently rapidly diversified in size and shape.
- *D. mauritiana* has smaller posterior lobes than *D. simulans* and more expansive expression of *Sox21b* in the developing genitalia. Using a reciprocal hemizygosity test, we show that variation in *Sox21b* underlies the diversification of epandrial posterior lobe size and shape between *D. simulans* and *D. mauritiana*.
- Behavioural tests show that the species allele of *Sox21b* causes differences in the duration of copulation in otherwise genetically identical backgrounds.
- *Sox21b* has evolved between *D. simulans* and *D. mauritiana*, and contributed to the divergence of a morphological novelty and copulatory behaviour between these two species.

## Results and Discussion

### Identification of transcription factors regulating the development of *Drosophila* male external genitalia

To better understand how the male genitalia develops and have evolved between *D. simulans* and *D. mauritiana* (Figure 1A), we used a candidate gene approach to interrogate RNA-seq data and genomic regions identified by introgression mapping^27^. We focussed on genes encoding transcription factors (TFs) because in many previous studies they have been shown to occupy key nodes in gene regulatory networks and contribute to morphological evolution^31–36^.

**Figure 1.**
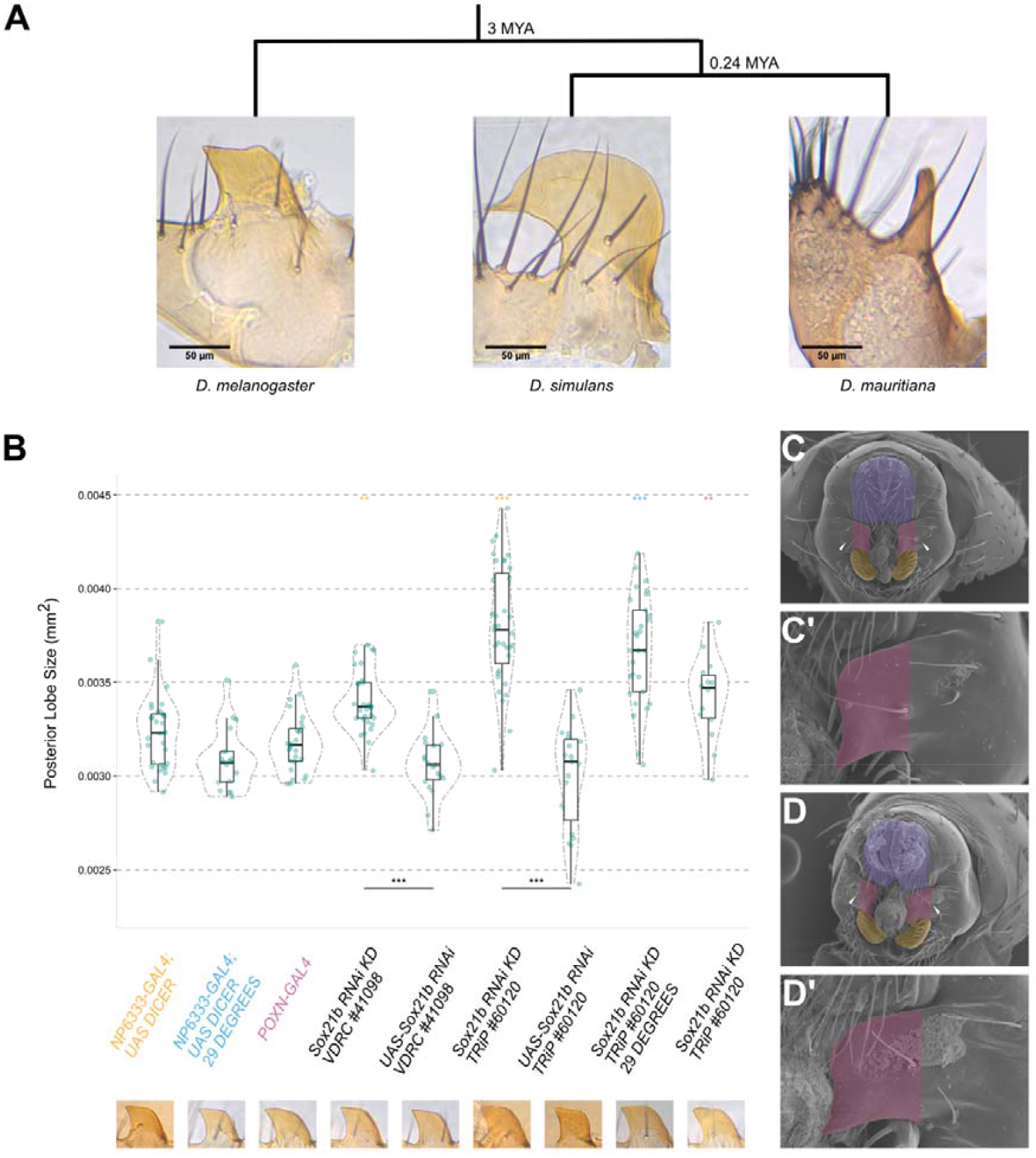
Effects on posterior lobe size from RNAi knockdown of *Sox21b* in *D. melanogaster*. (**A**) Posterior lobe of *D. melanogaster, D. simulans* and *D. mauritiana*. (**B**) RNAi knockdown of *Sox21b* in *D. melanogaster*. Asterisks above the knockdown line indicates the levels of statistical significance between the RNAi knockdown and associated parental GAL4 driver: red for *NP6333-GAL4*, blue for *NP6333-GAL4* at 29°C, and red for *Poxn-GAL4*. The lines below, adjoining the RNAi knockdown to the parental UAS-RNAi line, indicate the level of statistical significance between each pair. *** p < 0.001, and ** p < 0.01. Within each violin plot, the following values are represented: the median as the bold horizontal line, the box as the interquartile range, and the range as the vertical line. N > 13 for each line (Tables S1 and S2). (**C, C’, D, D’**) SEM images of example *NP6333-GAL4* control genitalia (**C, C**’) and RNAi genitalia after RNAi knockdown of *Sox21b* (**D, D’**). Purple highlights the cercus, pink the epandrial posterior lobes and yellow the surstyli.

Our previous analysis of RNA-Seq data from the developing male genitalia of *D. simulans* and *D. mauritiana* revealed 49 differentially expressed TF encoding genes^27^, 24 on the chromosome 3L arm, where we have previously generated high-resolution introgression maps of regions contributing to divergence in genital structures between these two species^26,27,37^. We tested the function of these 24 genes during genital development using RNAi knockdown in *D. melanogaster*. RNAi against ten of these genes significantly altered the size and/or bristle count of the posterior lobe, surstyli and/or cercus compared to parental controls (Table 1). Seven out of the ten affected just one structure, while *MED24, tna* and *E(spl)m3-HLH* knockdown affected multiple structures (Table 1 and Figure S1A-D). Only RNAi against *CKIIalpha-i1, knrl, MED10* (all cercus area) and *Sox21b* (posterior lobe area) altered structures consistent with the direction of the difference in their expression and phenotype in *D. mauritiana D1* compared to *D. simulans w*^*501*^ (Table 1; Figure S1A and S1C).

**Table 1.** Effects of RNAi knockdown on genital structures in *D. melanogaster* of previously identified genes encoding TFs on chromosome 3L. All RNAi knockdowns were conducted at 25°C. Effect sizes are denoted as ‘+/- (+/-)’ where +/- represents the effect size of the RNAi knockdown compared to the *NP6333-GAL4* control, and (+/-) to the *UAS-RNAi* control. * = <0.05, ** = <0.01, *** = <0.001. NS: Non-Significant. Gene names in bold font indicate more highly expressed in *D. mauritiana D1* and non-bold font more highly expressed in *D. simulans*^*w501*. X^Genes within an evolved region underlying divergence in posterior lobe morphology between *D. simulans* and *D. mauritiana* identified by previous introgression mapping^27^.

| Effects of RNAi Knockdown of Candidate Genes in *D. melanogaster* |  |  |  |  |
| --- | --- | --- | --- | --- |
| Gene | Epandrial<br>Posterior Lobe<br>Area | Cercus Area | Cercus Bristles | Surstylus<br>Bristles |
| **Asciz** | NS | +1.28***<br>(+1.04)** | NS | NS |
| **MED24** | -1.60***<br>(-0.78)* | NS | -2.09***<br>(-1.51)*** | -2.45***<br>(-1.89)*** |
| **tna** | -1.42***<br>(-0.93)* | NS | NS | +1.06**<br>(+1.37)*** |
| **CG17359<sup>X</sup>** | NS | NS | NS | +1.06**<br>(+1.37)*** |
| **Sox21b<sup>X</sup>** | +2.16***<br>(+2.63)*** | NS | NS | NS |
| *MED10* | NS | +0.81*<br>(+0.92)* | NS | NS |
| **Su(z)12** | NS | NS | +1.00**<br>(+0.98)* | NS |
| *knrl* | NS | +1.83***<br>(+2.09)*** | NS | NS |
| **CKIIalpha-i1** | NS | -1.13**<br>(-0.75)* | NS | NS |
| **E(spl)m3-HLH** | +1.11**<br>(+1.15)** | +1.01**<br>(+0.86)* | NS | NS |

From our RNAi knockdown experiments in *D. melanogaster, Sox21b* was the only differentially expressed gene between *D. mauritiana* and *D. simulans* within a mapped introgressed region to significantly affect the genitalia morphology consistent with the direction of the gene’s expression and phenotypic difference between these two species (Table 1)^27^. The posterior lobes were significantly larger when *Sox21b* expression was reduced by RNAi in *D. melanogaster* (with and without accounting for body size) evidencing a possible repressive function consistent with the expression and difference in size of this structure between *D. mauritiana* and *D. simulans* (Figure 1B, 1C and 1D; Table S1). This reveals a new role for *Sox21b* in male *Drosophila* genital development^38,39^. Furthermore, although the posterior lobes of the *D. melanogaster* species subgroup have been shown to have evolved from co-option of the Hox regulated gene network ancestrally involved in posterior spiracle formation^10^, none of the genes involved in subsequent posterior lobe diversification among these species have yet been identified. Therefore, we further examined the role of *Sox21b* in posterior lobe development and evolution.

### *Sox21b* is essential for the regulation of posterior lobe size

We first carried out additional RNAi knockdowns of *Sox21b* in *D. melanogaster*. We used a second UAS-RNAi line designed to target a different region of the *Sox21b* mRNA, an alternative driver line, *PoxN-GAL4*, (a posterior lobe specific GAL4 driver^40^), and increased the temperature to 29°C (Figure 1B). All the RNAi knockdowns of *Sox21b* resulted in an increase in posterior lobe area relative to controls (Figure 1B, 1C and 1D). In addition, the width of the base of the posterior lobes also significantly increased upon reduction of *Sox21b* expression (Figure S1E). Conversely, *Sox21b* RNAi led to a decrease in the size of the lateral plates i.e., the structure from which the posterior lobes grow out from (Figure S1F). The reduction of the lateral plate and reciprocal enlargement of the lobe reveals a potential trade-off in the proportion of cells assigned to posterior lobe versus lateral plate fate.

### Spatial differences in *Sox21b* expression between species in the posterior lobe primordium

Previous analysis of *Sox21b* in the developing male terminalia of *D. melanogaster* revealed expression in the developing posterior lobes and lateral plates during pupal stages^41^. We found that *Sox21b* is also expressed much earlier in the genital discs of *D. melanogaster, D. simulans* and *D. mauritiana* larvae (Figure 2).

**Figure 2:**
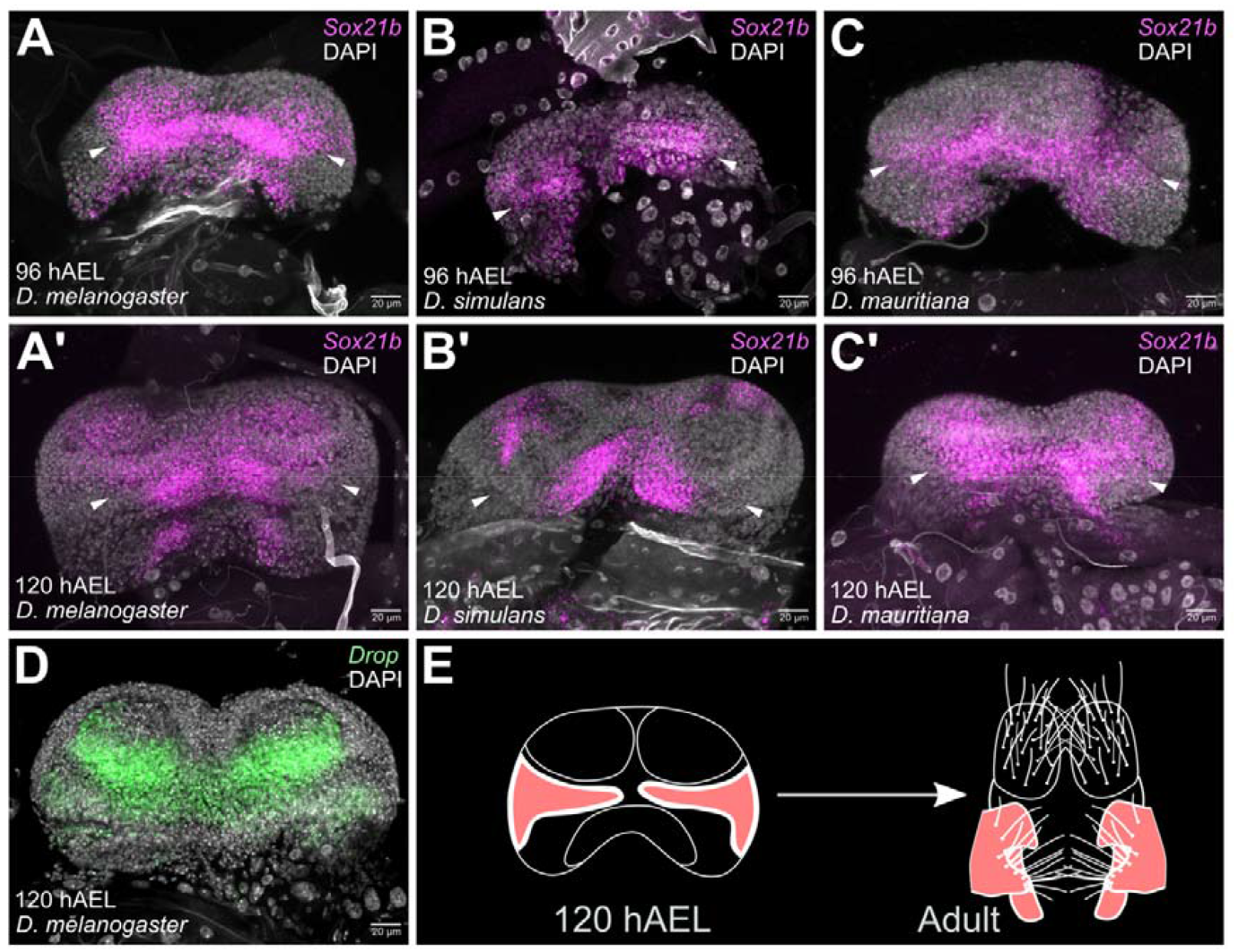
Dynamic developmental and evolutionary expression of *Sox21b* in *D. melanogaster, D. simulans* and *D. mauritiana*. (**A-C**) *Sox21b* Hybridisation Chain Reaction in situ hybridisation (HCR-ISH) of male genital discs at 96 hAEL and (**A’-C’**) 120 hAEL. (**A, A’**) *D. melanogaster*^*w1118*^. (**B, B’**) *D. simulans*^*w501*^. (**C, C’**) *D. mauritiana D1*. **A** (n=5), **A’** (n=7), **B** (n=3), **B’** (n=4), **C** (n=4), **C’** (n=4). (**D**) Anti-Drop staining in a *D. melanogaster*^*w1118*^ male genital disc (n=8). (**E**) Schematic showing origin of lateral plate, posterior lobe, and clasper primordium in *D. melanogaster*.

At 96 hours after egg laying (hAEL), *Sox21b* expression stretches across the medial and lateral regions of the genital discs of all three species, which represents the posterior lobe/lateral plate/clasper primordium (Figure 2A-C)^42^. However, in *D. mauritiana* expression was observed across the entirety of this area, whereas *D. simulans* lacks expression in the most lateral parts (Figures 2B and 2C). At 120 hAEL the medial expression contracts to varying extents among the species (Figure 2A’-C’). *D. simulans* exhibits the most extreme constriction in expression, with *Sox21b* mostly absent from the posterior lobe primordium (Figure 2B’), whereas in *D. mauritiana* the broader *Sox21b* expression persists (Figure 2C’). The expression of *Sox21b* in genital discs is similar to that of Drop (Figure 2D), which also regulates posterior lobe development^43^ and may even interact with Sox21b^44^.

Given our finding that *Sox21b* negatively regulates posterior lobe size, these data suggest that the higher and more expansive expression, detected by RNA-seq and in situ hybridisation respectively, in *D. mauritiana* compared to *D. simulans* contributes to the evolutionary difference in posterior lobe size between these two species.

### *Sox21b* causes the evolution of posterior lobe size between *D. simulans* and *D. mauritiana*

Since *Sox21b* regulates posterior lobe development in *D. melanogaster*, is located in a genomic region^27^ that contributes to differences in posterior lobe size between *D. mauritiana* and *D. simulans*, and is expressed differently between these two species, we then directly tested if this gene has contributed to the evolution of their posterior lobe size difference. To do this, we carried out a reciprocal hemizygosity test^37,45–47^. We used CRISPR/Cas9 to direct the insertion of 3XP3-DsRed fluorophore into exon 1 of *Sox21b* in a *D. simulans* line and an introgression line (IL108) carrying 2.8 Mb of *D. mauritiana* DNA spanning region P5^27^ in an otherwise *D. simulans* genome (Figure S2A). This successfully disrupted the reading frame in the *Sox21b* locus from both species (Figure S2A). Interestingly, *D. simulans Sox21b* mutants were homozygous viable, and these flies had significantly larger posterior lobes than controls, consistent with *Sox21b* RNAi knockdown in *D. melanogaster* (Tables S2 and S3 Figure 1B and S2B).

Reciprocal hemizygotes were then generated by crossing the two sets of independent *Sox21b* mutant lines to generate flies that were genetically identical except they had either a working *D. mauritiana Sox21b* allele or a working *D. simulans Sox21b* allele (Figure 3A). Remarkably, posterior lobe size was significantly different depending upon the species-origin of the working *Sox21b* allele and consistent with the direction of the species difference (Figure 3A). Male flies with the *D. mauritiana Sox21b* allele had significantly smaller lobes with narrower bases compared to flies with a working *D. simulans Sox21b allele* (Cohens’ effect size = − 1.172, Figures 3A and S2C). Surstylus bristle count, cercus bristle count and cercus area did not differ significantly between the reciprocal hemizygotes (Tables S2 and S3). These results show that variation in *Sox21b* has contributed to the evolution of posterior lobe size between *D. mauritiana* and *D. simulans*.

**Figure 3:**
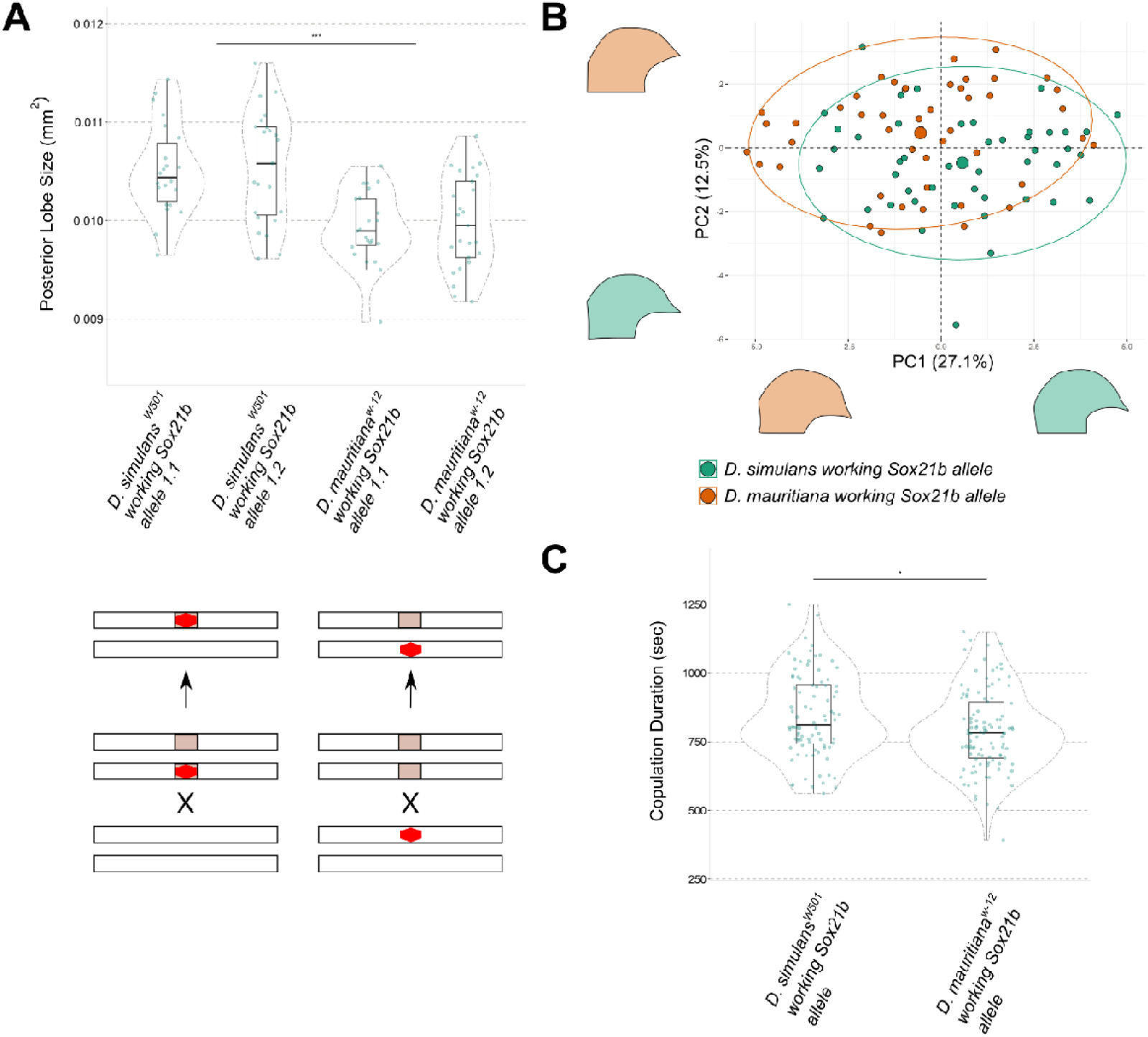
Effects of species-specific *Sox21b* alleles on posterior lobe size and shape. (**A**) Posterior lobes are smaller in size when only the *D. mauritiana* allele of *Sox21b* is working (n > 22). (Tables S2 and S3). The lower schematic illustrates the generation of reciprocal hemizygotes containing either a *D. simulans* working allele or *D. mauritiana* working allele of *Sox21b*. The red shape represents the disrupted *Sox21b* locus, the shaded rectangle represents the *D. mauritiana* introgressed region. (**B**) PC1 versus PC2, whereby both statistical identifiers summarise 37.6% of total shape variation between the two reciprocal hemizygotes. Outlines of lobes show the minimal and maximal data points for each principal component. (**C**) The *D. mauritiana* working *Sox21b* allele significantly decreased copulation duration compared to the working *D. simulans Sox21b* allele (n > 87 for each reciprocal hemizygote) (Table S3).

### The shape of the epandrial posterior lobe is altered according to the species origin of *Sox21b*

We next tested if variation in *Sox21b* contributes to the evolution of posterior lobe shape between *D. simulans* and *D. mauritiana*. Using Elliptical Fourier Analysis (EFA), we summarised the morphometric changes in shape between the reciprocal hemizygotes to identify variation originating from species-specific alleles of *Sox21b* (Figures 2B, S2D and S2E; Table S4). The two PCs summarising the highest proportion of shape variation, PC1 and PC2 contributing 27.1% and 12.5% respectively, were significantly different between the two reciprocal hemizygotes (Figures S2E and S2E’). When assessing the significant variation captured by PC2, a notable shape alteration in the ‘beak’ region of the lobe (extending from the main body of the lobe, parallel to the artificial baseline) was identified (Figure 3B). PC1 captured shape alterations from the artificial baseline to the beginning of the beak extension. The *D. mauritiana Sox21b* allele reduced the height of the ‘neck’ leading up to the ‘beak’ extension, in comparison to *D. simulans* working allele (Figure 3B). Therefore, EFA of the posterior lobes between the *Sox21b* reciprocal hemizygotes revealed shape variation affecting the beak extension and the neck consistent with the direction of the species difference.

### The behavioural consequences of *Sox21b* species-specific alleles

Finally we sought to understand if the species-specific alleles of *Sox21b* that contribute to evolutionary differences in posterior lobe morphology also cause detectable differences in copulation by crossing individually reciprocal hemizygote males to *D. simulans*^*w501*^ females. In contrast to a previous study that used laser microdissection to alter the morphology of the posterior lobes in *D. simulans*^16^, we found no difference in mating frequency between reciprocal hemizygote males (Tables S3 and S5). We also found no difference in copulation latency between reciprocal hemizygote males (Tables S3 and S5) but indeed, to the best of our knowledge, copulation latency differences do not segregate in these two species, although a previous study using introgression lines between *D. mauritiana* and *D. sechellia* found that PL morphology did affect copulation latency^14^. Interestingly, however, we did observe that males carrying a working *D. mauritiana Sox21b* allele engaged in significantly shorter copulations than those carrying a working *D. simulans Sox21b* allele, which may result in a difference in sperm transfer time and could likely affect fitness (Figure 3C; Table S3). This result is consistent with the previous study of *D. mauritiana* and *D. sechellia*, and more importantly a species-specific difference in copulation duration, with shorter copulations in *D. mauritiana* than in *D. simulans*^48–50^.

## Conclusions

We have shown that inter-specific allelic variation in *Sox21b* contributes to the diversification of an evolutionary novelty, the posterior lobes, and this affects the copulatory behaviour suggesting this gene has been subject to sexual selection to help shape lobes with different morphologies. Genes have been identified that contribute to the evolution of other sexual traits^37,51,52^, but *Sox21b* is the first for the diversification of the posterior lobes to the best of our knowledge. It appears that *Sox21b* regulates the early stages of posterior lobe development and allocates which cells from the primordial lateral plate will contribute to this structure by restricting the allocation of cells to posterior lobe fate and influencing the size and shape of these structures. Our results strongly suggest a difference in *Sox21b* expression between *D. mauritiana* and *D. simulans* underlies the role of this gene in the evolution of posterior lobe morphology between these two species, but we cannot rule out that coding changes may also be involved. It will be important to identify the other genes that contribute to posterior lobe diversity between *D. mauritiana* and *D. simulans*, and other species, to understand more fully how the gene regulatory network for posterior lobes was co-opted and evolved. Several other genes have already been found that regulate posterior lobe development^10,28,41^ and a crucial role has been identified for the apical ECM in cell extension from the lateral plate^53^. Therefore it will be interesting to determine how *Sox21b* is integrated into this network.

## Resource availability

### Lead contact

Further information and requests for resources and reagents should be directed to and will be fulfilled by the lead contact Alistair McGregor

### Materials availability

All unique/stable reagents generated in this study are available from the lead contact without restriction.

### Data and code availability

RNA-seq data underlying this study are deposited in the ArrayExpress database at EMBL-EBI under accession number E-MTAB-9465 (https://www.ebi.ac.uk/arrayexpress/experiments/E-MTAB-9465).

## Materials and Methods

### RNAi knockdown of differentially expressed transcription factors in *D. melanogaster*

Differentially expressed TFs were identified from RNA-Seq data generated from the developing male genitalia of *D. simulans* and *D. mauritiana* at 30 – 36 hours after puparium formation as previously reported^27^. Those selected for further analysis in this study were filtered based on chromosome 3 genomic location with respect to previous QTL and introgression mapping studies^26,27,30,37^. Selected differentially expressed TFs were assessed for roles in the development of the male periphallic genitalia using the GAL4-UAS system to drive RNAi in *D. melanogaster*. UAS-RNAi lines for each gene were provided by the Vienna Drosophila Resource Centre (VDRC)^54^ and from Bloomington Drosophila Stock Centre (BDSC) (NIH P40OD018537). UAS-RNAi males were crossed to *NP6333-GAL4 (P(GawB)PenNP6333)* virgin females, (which drives GAL4 expression in all imaginal discs from larval stages and during metamorphosis) also carrying *UAS-Dicer-2 (P[UAS-Dcr-2*.*D]*^55^. *Sox21b* RNAi was repeated using the *Poxn-GAL4 (14*.*1*.*1)* driver, which specifically drives in the posterior lobe primordium from larval imaginal disc stage^40^ (Table S1). All crosses were carried out at 25ºC, with selected crosses repeated at 29ºC to enhance the GAL4^56^ (Table S1). All crosses were performed using a 1:2 male to female ratio. The same conditions were used for each control line, as well as reciprocal hemizygote crosses. Three biological replicates, with a total sample size of n > 14, were phenotyped for each cross. Crosses were transferred to standard cornmeal food every two days and maintained in a 12-hour light/dark cycle incubators. Males aged at least 3 days were collected and then stored in 70% EtOH at −20ºC for phenotyping.

### Phenotyping of RNAi knockdown flies

The cercus, epandrial posterior lobes and surstyli of the adult male genitalia were dissected in Hoyer’s Solution using 0.14 mm diameter stainless steel pins and then mounted in Hoyer’s solution. This was done using slides containing eight individual 6 mm diameter wells. To account for body size, the T2 legs of each fly were also dissected and mounted. A Zeiss Axioplan light microscope with a Jenoptik ProgRes C3 camera was used to image each dissected structure. 250X magnification was used for the genital structures, and 160X magnification for the T2 legs. The area of posterior lobes and cercus, and length of T2 tibias were measured manually using ImageJ^57^ (Table S2). The bristles were counted using the light microscope and a tap counter. When drawing the outline of the posterior lobe, an artificial baseline was used as previously described^28^. The area and bristle count were recorded for both pairs of structures per individual and the average was then used for the latter statistical analysis.

### Generation of Sox21b-DsRed

To disrupt the reading frame of *Sox21b*, 3XP3-DsRed was inserted 152 bp into exon 1 using CRISPR/Cas9^58^ (Figure 2C). This exon was chosen because it did not include restriction sites required in the cloning procedure 1 kb either side from the gRNA cut site, as well as bypassing the conserved HMG-Box domain that may have resulted in off-target effects. The gRNAs were designed using FlyCRISPR^59^ and inserted into the pCFD3 plasmid^58^. The homology arms (HA) were amplified by PCR from salt extracted genomic DNA of the focal strains (adapted from^60^), and inserted into the plasmid pHD-DsRed^59^. Plasmids were sequenced prior to injections to verify homology arm and gRNA incorporation. 200 ng/μl of gRNA-pCFD3 and 500 ng/μl of HA-pHD-DsRed were injected (using a Eppendorf FemtoJet 4i and Leica light microscope) into *D. simulans*^*w501*^ and *IL108* embryos^27^, both carrying *nanos-Cas9* on the X chromosome. 48 hours prior to injection, cages were set up containing apple juice plate and yeast paste, which were changed twice per day prior to injections. Surviving adults from injected embryos were then backcrossed to non-injected adults of the same strain. Progeny were screened for the DsRed marker in their eyes using a Zeiss Axiozoom microscope and those positive were amplified and sequenced to verify genome editing (Figure 2C).

### Generation of reciprocal hemizygotes

To generate the reciprocal hemizygote males, transgenic *D. simulans*^*w501*^ and *IL108* males carrying *Sox21b-DsRed* were crossed to *IL108* and *D. simulans*^*w501*^ virgin females, respectively. F1 males with *Sox21b-DsRed* were kept for phenotypic analysis of their posterior lobes (Tables S2 and S3).

### Scanning electron microscopy

Flies stored in 70% EtOH were moved to 100% EtOH at least 24 hours prior to imaging. The posterior of the fly was dissected in EtOH. Samples were processed in a critical point dryer and mounted on SEM stubs, then gold coated for 30 seconds. The genitalia were imaged using SE mode at 5 kV in a Hitachi S-3400N SEM, with a working distance of 13 to 14 mm. Whole genitalia was imaged at a magnification of 250x, and individual periphallic structures at 900x.

### Posterior lobe shape analysis

Posterior lobes were manually traced as described above. To quantitatively assess shape variation, PAT-GEOM^61^ was used to perform Elliptical Fourier Analysis (EFA) on each region of interest (ROI)^62^. This software benefitted from being trace-start point, scale, rotation, and translation insensitive. One posterior lobe at random was assessed per fly (n = 45), and twenty descriptors were assigned to each ROI for EFA (Table S4). Principal component analysis (PCA) was performed using prcomp and factoextra^63^ in R, to evaluate variation between lines. This package standardised the data to have a mean of zero and variance of one prior to computing the PCA. The eigenvalues for each principal component were also computed to identify those with a value above 1, to which these principal components were retained for analysis (Figure S2D). Outlines of lobes corresponding to the extremities of the minimum and maximum values of the principal components labelled on each axis were extracted from the ROI data.

### HCR-ISH of *Sox21b* in genital discs

To capture *Sox21b* expression, we carried out HCR-ISH on genital discs from L2 (96 hAEL) and L3 larvae (120 hAEL) to complement previous ISH data of *Sox21b* in developing male terminalia that showed expression in the posterior lobes and lateral plates^41^. *D. melanogaster*^*w1118*,^ *D. simulans*^*w501*^ and *D. mauritiana D1* larvae were dissected in ice cold 1XPBS. Each larva was cut in half and the posterior half inverted then placed into 4% formaldehyde in 0.3% PBT (TRITON X-100) for 20 minutes. The HCR procedure was based on an established protocol^64^. The probe from Molecular Instruments to target *Sox21b* was designed as 20 individual hairpins spanning the entirety of the gene, ensuring that all isoforms were captured^65^. A 16 nM probe solution was used, and the sample was incubated for 24 hours at 37°C on an orbital shaker at 60 RPM. Note, we used half the volumes of each solution compared to the protocol^64^. DAPI was diluted in the probe wash solution for nuclear staining of the samples. Samples were then dissected using forceps, mounted in 1X PBS and imaged. Images were obtained on the Zeiss LSM800 upright Confocal Laser Scanning Microscope with a 20X objective.

### Immunohistochemistry in genital discs

Genital discs of L3 larvae from *D. melanogaster* were dissected and fixed as described for HCR-ISH. Samples were incubated in 10% NGS in 0.1% PBST (Tween-20) for 1 hour prior to the addition of the primary antibody. 1:200 dilution of Rabbit Anti-Drop was used, and samples were incubated overnight at 4 degrees. 1:600 Anti-Rabbit 488 secondary was incubated with sample the following day in 10% NGS at 4ºC overnight. Samples were then dissected and mounted in PBS, and imaged that same day with the same imaging parameters as described above. As *Drop* is a male-specifically expressed factor, female genital discs could be readily sorted from male discs when imaging^43^.

### Behaviour assays using *Sox21b* reciprocal hemizygotes

All mating assays were carried out at 25°C and 70% humidity. Flies were kept in these conditions 24 hours prior to their respective mating assay, allowing acclimatisation. Flies used in the mating assays were reared at 25°C in a 12-hour light/dark cycle. Mating experiments were carried out within the first hour of lights as previously described^66^. Males were collected as pupae (identified by the presence of sex combs) and aged individually for 4-5 days in separate vials, ensuring they were socially naive to account for the influence of male aggression and competition in the presence of other males. Females were similarly collected as pupae (identified by the absence of sex combs) and aged for 4-5 days in vials in groups of 5-10, ensuring virgin status. Reciprocal hemizygote males were screened for the DsRed marker at least 48 hours prior to the mating to ensure full recovery from the brief CO_2_ exposure. Single males were paired with single females in a standard food vial with the stopper pushed down to 1 cm above the food, creating a restricted ‘mating chamber’. Each pairing was observed for a total of 90 minutes. Mating frequency refers to whether there was evidence of mating in the 90-minute observation period (Y/N), characterised as mounting of the female by the male for a minimum of 7 minutes because this is considered the minimal time for sperm transfer to occur^19,48^. Copulation latency was measured as the time between pairing and copulation onset. Copulation duration was quantified as the time elapsed from initial male mounting and dismounting the female.

### Quantification and statistical analysis

All statistical analyses were carried out using R version 4.2.0^67^. Measurements of each structure described above were first assessed for normality using the Shapiro Wilk test. Normally distributed data were analysed using Dunnett’s test and ANOVA, whereas the Kruskal Wallis test was used for non-normally distributed data^68^. All comparisons included the RNAi knockdown compared to both parental controls. If this test was evaluated as statistically significant, the Tukey’s Test / Wilcoxon Rank Sum Test (BH p-adjusted method^69^) was used to identify if the RNAi knockdown was significantly different to both parental controls (Table S1). Results were concluded as non-significant when p =/> 0.05 or the effect detected in the RNAi knockdown was an intermediate of the two parental controls.

Where the T2 tibia length was significantly different to both parental controls following the tests described above, Pearson/Spearman analysis was performed on the cercus and posterior lobe area of these crosses dependent on Shapiro Wilk test results. If statistically significant, the area of the structure was divided by the tibia length squared, and statistical tests were carried out on the normalised version of the measurements (Table S1). Effect sizes between each parental control to the RNAi knockdown were calculated using Cohens’ d, where the coefficient value represents the number of standard deviations different between the two population means under investigation^70^. To calculate the Cohens’ d coefficient, the difference of the means from the two populations of samples were divided by the cumulative pooled standard deviation of them both.

The phenotypic measurements for the reciprocal hemizygotes and null mutants were analysed using an independent t-test or Wilcoxon Rank Sum Test dependent on the normality of the data. Details of morphometric analysis of posterior lobe shape can be found above. For PCA, the Wilcoxon Rank Sum Test was used.

A Fisher’s exact test was conducted to analyse copulation frequency. The Shapiro Wilk test was conducted to assess the distribution of the datasets. As the copulation latency dataset was not normally distributed, a Kruskal Wallis test was performed. Copulation duration was analysed using an independent t-test. Violin plots indicate the mean of the data, with the first quartile and the third quartile value shown. The width of the violin plots represents the frequency of data points at assigned values.

## Acknowledgements

This study was funded in part by BBSRC (BB/X006689/1) and NERC grants (NE/M001040/1) to APM and MDSN, a BBSRC DTP studentship (BB/M011224/1) to AR and an Oxford Brookes University Nigel Groome studentship to EH. We thank the Bloomington *Drosophila* Stock Center and Vienna *Drosophila* RNAi Center for fly lines, as well as Molecular Instruments for the custom made *Sox21b* HCR probe. We also thank members of the Rebeiz lab for their thoughtful discussions.

## Author contributions

The project was conceived by APM and MDSN. Experiments were carried out by AR and EH. Data was analysed by all authors. AR, APM and MDSN wrote the paper assisted by EH and JFJ.

## Supplemental information

Figures S1 to S3 and Tables S1 to S5 are provided as supplemental information.

## References

1. York, J.R., and McCauley, D.W. (2020). The origin and evolution of vertebrate neural crest cells. Open Biol 10, 190285. 10.1098/rsob.190285.

2. Tomoyasu, Y. (2021). What crustaceans can tell us about the evolution of insect wings and other morphologically novel structures. Curr Opin Genet Dev 69, 48–55. https://doi.org/10.1016/j.gde.2021.02.008.

3. Moczek, A.P. (2009). On the origins of novelty and diversity in development and evolution: a case study on beetle horns. In Cold Spring Harbor Symposia on Quantitative Biology (Citeseer), p. 289.

4. Kijimoto, T., Pespeni, M., Beckers, O., and Moczek, A.P. (2013). Beetle horns and horned beetles: emerging models in developmental evolution and ecology. WIREs Developmental Biology 2, 405–418. https://doi.org/10.1002/wdev.81.

5. Rebeiz, M., and Tsiantis, M. (2017). Enhancer evolution and the origins of morphological novelty. Curr Opin Genet Dev 45, 115–123. https://doi.org/10.1016/j.gde.2017.04.006.

6. Shubin, N., Tabin, C., and Carroll, S. (2009). Deep homology and the origins of evolutionary novelty. Nature 457, 818–823. 10.1038/nature07891.

7. Bruce, H.S., and Patel, N.H. (2022). The Daphnia carapace and other novel structures evolved via the cryptic persistence of serial homologs. Current Biology 32, 3792–3799.e3. https://doi.org/10.1016/j.cub.2022.06.073.

8. Colizzi, E.S., Hogeweg, P., and Vroomans, R.M.A. (2022). Modelling the evolution of novelty: a review. Essays Biochem 66, 727–735. 10.1042/EBC20220069.

9. Muller, G.B., and Wagner, G.P. (1991). Novelty in Evolution: Restructuring the Concept. Annu Rev Ecol Syst 22, 229–256. 10.1146/annurev.es.22.110191.001305.

10. Glassford, W.J., Johnson, W.C., Dall, N.R., Smith, S.J., Liu, Y., Boll, W., Noll, M., and Rebeiz, M. (2015). Co-option of an Ancestral Hox-Regulated Network Underlies a Recently Evolved Morphological Novelty. Dev Cell 34, 520–531. 10.1016/j.devcel.2015.08.005.

11. Jagadeeshan, S., and Singh, R.S. (2006). A time-sequence functional analysis of mating behaviour and genital coupling in Drosophila: role of cryptic female choice and male sex-drive in the evolution of male genitalia. J Evol Biol 19, 1058–1070. 10.1111/j.1420-9101.2006.01099.x.

12. Yassin, A., and Orgogozo, V. (2013). Coevolution between Male and Female Genitalia in the Drosophila melanogaster Species Subgroup. PLoS One 8, e57158. 10.1371/journal.pone.0057158.

13. Kopp, A., and True, J.R. (2002). Evolution of male sexual characters in the Oriental Drosophila melanogaster species group. Evol Dev 4, 278–291. https://doi.org/10.1046/j.1525-142X.2002.02017.x.

14. Frazee, S.R., Harper, A.R., Afkhami, M., Wood, M.L., McCrory, J.C., and Masly, J.P. (2021). Interspecific introgression reveals a role of male genital morphology during the evolution of reproductive isolation in Drosophila. Evolution (N Y) 75, 989–1002. 10.1111/evo.14169.

15. Frazee, S.R., and Masly, J.P. (2015). Multiple sexual selection pressures drive the rapid evolution of complex morphology in a male secondary genital structure. Ecol Evol 5, 4437–4450. 10.1002/ece3.1721.

16. LeVasseur-Viens, H., Polak, M., and Moehring, A.J. (2015). No evidence for external genital morphology affecting cryptic female choice and reproductive isolation in Drosophila. Evolution (N Y) 69, 1797–1807. 10.1111/evo.12685.

17. Robertson, H.M. (1988). Mating asymmetries and phylogeny in the Drosophila melanogaster species complex.

18. McDermott, S.R., and Kliman, R.M. (2008). Estimation of Isolation Times of the Island Species in the Drosophila simulans Complex from Multilocus DNA Sequence Data. PLoS One 3, e2442..

19. Coyne, J.A. (1983). Genetic Basis of Differences in Genital Morphology Among Three Sibling Species of Drosophila. Evolution (N Y) 37, 1101–1118. 10.2307/2408834.

20. House, C.M., Lewis, Z., Hodgson, D.J., Wedell, N., Sharma, M.D., Hunt, J., and Hosken, D.J. (2013). Sexual and Natural Selection Both Influence Male Genital Evolution. PLoS One 8, e63807. 10.1371/JOURNAL.PONE.0063807.

21. House, C.M., Lewis, Z., Sharma, M.D., Hodgson, D.J., Hunt, J., Wedell, N., and Hosken, D.J. (2021). Sexual selection on the genital lobes of male Drosophila simulans. Evolution (N Y) 75, 501–514. 10.1111/EVO.14158.

22. Liu, J., Mercer, J.M., Stam, L.F., Gibson, G.C., Zengt, Z.-B., and Laurie, C.C. (1996). Genetic Analysis of a Morphological Shape Difference in the Male Genitalia of Drosophila simulans and D. mauritiana. Genetics 142, 1129–1145. 10.1093/genetics/142.4.1129.

23. Zeng, Z.B., Liu, J., Stam, L.F., Kao, C.H., Mercer, J.M., and Laurie, C.C. (2000). Genetic architecture of a morphological shape difference between two Drosophila species. Genetics 154, 299–310.

24. LeVasseur-Viens, H., and Moehring, A.J. (2014). Individual Genetic Contributions to Genital Shape Variation between Drosophila simulans and D. mauritiana. Int J Evol Biol 2014, 1–9. 10.1155/2014/808247.

25. True, J.R., Liu, J., Stam, L.F., Zeng, Z.-B., and Laurie, C.C. (1997). Quantitative Genetic Analysis of Divergence in Male Secondary Sexual Traits Between Drosophila simulans and Drosophila mauritiana. Evolution (N Y) 51, 816–832. 10.2307/2411157.

26. Tanaka, K.M., Hopfen, C., Herbert, M.R., Schlötterer, C., Stern, D.L., Masly, J.P., McGregor, A.P., and Nunes, M.D.S. (2015). Genetic Architecture and Functional Characterization of Genes Underlying the Rapid Diversification of Male External Genitalia Between Drosophila simulans and Drosophila mauritiana. Genetics 200, 357. 10.1534/genetics.114.174045.

27. Hagen, J.F.D., Mendes, C.C., Booth, S.R., Figueras Jimenez, J., Tanaka, K.M., Franke, F.A., Baudouin-Gonzalez, L., Ridgway, A.M., Arif, S., Nunes, M.D.S., et al. (2021). Unraveling the Genetic Basis for the Rapid Diversification of Male Genitalia between Drosophila Species. Mol Biol Evol 38, 437–448. 10.1093/MOLBEV/MSAA232.

28. Hackett, J.L., Wang, X., Smith, B.R., and Macdonald, S.J. (2016). Mapping QTL Contributing to Variation in Posterior Lobe Morphology between Strains of Drosophila melanogaster. PLoS One 11, e0162573–e0162573. 10.1371/journal.pone.0162573.

29. Laurie, C.C., True, J.R., Liu, J., and Mercer, J.M. An Introgression Analysis of Quantitative Trait Loci That Contribute to a Morphological Difference Between Drosophila simulans and D. mauritiana.

30. Masly, J.P., Dalton, J.E., Srivastava, S., Chen, L., and Arbeitman, M.N. (2011). The Genetic Basis of Rapidly Evolving Male Genital Morphology in Drosophila. Genetics 189, 357. 10.1534/genetics.111.130815.

31. Zhang, L., Mazo-Vargas, A., and Reed, R.D. (2017). Single master regulatory gene coordinates the evolution and development of butterfly color and iridescence. Proceedings of the National Academy of Sciences 114, 10707–10712. 10.1073/pnas.1709058114.

32. Williams, T.M., Selegue, J.E., Werner, T., Gompel, N., Kopp, A., and Carroll, S.B. (2008). The Regulation and Evolution of a Genetic Switch Controlling Sexually Dimorphic Traits in Drosophila. Cell 134, 610–623. 10.1016/j.cell.2008.06.052.

33. Stern, D.L. (2011). Evolution, Development, and the Predictable Genome.

34. Chan, Y.F., Marks, M.E., Jones, F.C., Villarreal, G., Shapiro, M.D., Brady, S.D., Southwick, A.M., Absher, D.M., Grimwood, J., Schmutz, J., et al. (2010). Adaptive Evolution of Pelvic Reduction in Sticklebacks by Recurrent Deletion of a Pitx1 Enhancer. Science (1979) 327, 302–305. 10.1126/science.1182213.

35. Stern, D.L., and Franke, N. (2013). The structure and evolution of cis-regulatory regions: the shavenbaby story. Philosophical Transactions of the Royal Society B: Biological Sciences 368. 10.1098/RSTB.2013.0028.

36. Carroll, S.B., Grenier, J.K., and Weatherbee, S.D. (2009). From DNA to Diversity: Molecular Genetics and the Evolution of Animal Design 2nd ed. (Wiley-Blackwell).

37. Hagen, J.F.D., Mendes, C.C., Blogg, A., Payne, A., Tanaka, K.M., Gaspar, P., Figueras Jimenez, J., Kittelmann, M., McGregor, A.P., and Nunes, M.D.S. (2019). tartan underlies the evolution of Drosophila male genital morphology. Proceedings of the National Academy of Sciences 116, 19025. 10.1073/pnas.1909829116.

38. McKimmie, C., Woerfel, G., and Russell, S. (2005). Conserved genomic organisation of Group B Sox genes in insects. BMC Genet 6, 26. 10.1186/1471-2156-6-26.

39. Akhund-Zade, J., Lall, S., Gajda, E., Yoon, D., Ayroles, J.F., and de Bivort, B.L. (2021). Genetic basis of offspring number-body weight tradeoff in Drosophila melanogaster. G3 (Bethesda) 11, jkab129. 10.1093/g3journal/jkab129.

40. Boll, W., and Noll, M. (2002). The Drosophila Pox neuro gene: control of male courtship behavior and fertility as revealed by a complete dissection of all enhancers. Development 129, 5667–5681. 10.1242/dev.00157.

41. Vincent, B.J., Rice, G.R., Wong, G.M., Glassford, W.J., Downs, K.I., Shastay, J.L., Charles-Obi, K., Natarajan, M., Gogol, M., Zeitlinger, J., et al. (2019). An Atlas of Transcription Factors Expressed in Male Pupal Terminalia of Drosophila melanogaster. G3 Genes|Genomes|Genetics 9, 3961–3972. 10.1534/g3.119.400788.

42. Keisman, E.L., and Baker, B.S. (2001). The Drosophila sex determination hierarchy modulates wingless and decapentaplegic signaling to deploy dachshund sex-specifically in the genital imaginal disc. Development 128, 1643–1656. 10.1242/dev.128.9.1643.

43. Chatterjee, S.S., Uppendahl, L.D., Chowdhury, M.A., Ip, P.-L., and Siegal, M.L. (2011). The female-specific Doublesex isoform regulates pleiotropic transcription factors to pattern genital development in Drosophila. Development 138, 1099. 10.1242/dev.055731.

44. Shokri, L., Inukai, S., Hafner, A., Basler, K., Deplancke, B., and Bulyk, M.L. (2019). A Comprehensive Drosophila melanogaster Transcription Factor Interactome. Cell Rep 27, 955–970. https://doi.org/10.1016/j.celrep.2019.03.071.

45. Stern, D.L. (2014). Identification of loci that cause phenotypic variation in diverse species with the reciprocal hemizygosity test. Trends in Genetics 30, 547–554. https://doi.org/10.1016/j.tig.2014.09.006.

46. Lamb, A.M., Wang, Z., Simmer, P., Chung, H., and Wittkopp, P.J. (2020). ebony Affects Pigmentation Divergence and Cuticular Hydrocarbons in Drosophila americana and D. novamexicana. Front Ecol Evol 8.

47. Ding, Y., Berrocal, A., Morita, T., Longden, K.D., and Stern, D.L. (2016). Natural courtship song variation caused by an intronic retroelement in an ion channel gene. Nature 536, 329–332. 10.1038/nature19093.

48. Price, C.S.C., Kim, C.H., Gronlund, C.J., and Coyne, J.A. (2001). CRYPTIC REPRODUCTIVE ISOLATION IN THE DROSOPHILA SIMULANS SPECIES COMPLEX. Evolution (N Y) 55, 81–92. 10.1111/j.0014-3820.2001.tb01274.x.

49. Cobb, M., Burnet, B., and Connolly, K. (1988). Sexual isolation and courtship behavior in Drosophila simulans, D. mauritiana, and their interspecific hybrids. Behav Genet 18, 211–225. 10.1007/BF01067843.

50. Coyne, J.A. (1993). THE GENETICS OF AN ISOLATING MECHANISM BETWEEN TWO SIBLING SPECIES OF DROSOPHILA. Evolution (N Y) 47, 778–788. 10.1111/j.1558-5646.1993.tb01233.x.

51. Nagy, O., Nuez, I., Savisaar, R., Peluffo, A.E., Yassin, A., Lang, M., Stern, D.L., Matute, D.R., David, J.R., and Courtier-Orgogozo, V. (2018). Correlated Evolution of Two Copulatory Organs via a Single <em>cis</em>-Regulatory Nucleotide Change. Current Biology 28, 3450–3457.e13. 10.1016/j.cub.2018.08.047.

52. Gao, J., Barmina, O., Thompson, A., Kim, B.Y., Suvorov, A., Tanaka, K., Watabe, H., Toda, M.J., Chen, J.-M., Katoh, T.K., et al. (2022). Secondary reversion to sexual monomorphism associated with tissue-specific loss of doublesex expression. Evolution (N Y) 76, 2089–2104. 10.1111/evo.14564.

53. Smith, S.J., Davidson, L.A., and Rebeiz, M. (2020). Evolutionary expansion of apical extracellular matrix is required for the elongation of cells in a novel structure. Elife 9, e55965. 10.7554/eLife.55965.

54. Dietzl, G., Chen, D., Schnorrer, F., Su, K.-C., Barinova, Y., Fellner, M., Gasser, B., Kinsey, K., Oppel, S., Scheiblauer, S., et al. (2007). A genome-wide transgenic RNAi library for conditional gene inactivation in Drosophila. Nature 448, 151–156. 10.1038/nature05954.

55. Stieper, B.C., Kupershtok, M., Driscoll, M. V., and Shingleton, A.W. (2008). Imaginal discs regulate developmental timing in Drosophila melanogaster. Dev Biol 321, 18–26. 10.1016/J.YDBIO.2008.05.556.

56. Blake, A.J., Finger, D.S., Hardy, V.L., and Ables, E.T. (2017). RNAi-Based Techniques for the Analysis of Gene Function in Drosophila Germline Stem Cells. In RNAi and Small Regulatory RNAs in Stem Cells: Methods and Protocols, B. Zhang, ed. (Springer New York), pp. 161–184. 10.1007/978-1-4939-7108-4_13.

57. Schneider, C.A., Rasband, W.S., and Eliceiri, K.W. (2012). NIH Image to ImageJ: 25 years of image analysis. Nat Methods 9, 671–675. 10.1038/nmeth.2089.

58. Gratz, S.J., Rubinstein, C.D., Harrison, M.M., Wildonger, J., and O’Connor-Giles, K.M. (2015). CRISPR-Cas9 Genome Editing in Drosophila. Curr Protoc Mol Biol 111, 31.2.1–31.2.20. 10.1002/0471142727.mb3102s111.

59. Gratz, S.J., Ukken, F.P., Rubinstein, C.D., Thiede, G., Donohue, L.K., Cummings, A.M., and O’Connor-Giles, K.M. (2014). Highly Specific and Efficient CRISPR/Cas9-Catalyzed Homology-Directed Repair in Drosophila. Genetics 196, 961–971. 10.1534/genetics.113.160713.

60. Miller, S.A., Dykes, D.D., and Polesky, H.F. (1988). A simple salting out procedure for extracting DNA from human nucleated cells. Nucleic Acids Res 16, 1215. 10.1093/nar/16.3.1215.

61. Chan, I., Stevens, M., and Todd, P. (2018). PAT-GEOM: A Software Package for the Analysis of Animal Patterns. Methods Ecol Evol 10. 10.1111/2041-210x.13131.

62. Caple, J., Byrd, J., and Stephan, C.N. (2017). Elliptical Fourier analysis: fundamentals, applications, and value for forensic anthropology. Int J Legal Med 131, 1675–1690. 10.1007/s00414-017-1555-0.

63. Kassambara, A., and Mundt, F. (2020). Extract and Visualize the Results of Multivariate Data Analyses.

64. Younger, M.A., Herre, M., Goldman, O. V, Lu, T.-C., Caballero-Vidal, G., Qi, Y., Gilbert, Z.N., Gong, Z., Morita, T., Rahiel, S., et al. (2022). Non-Canonical Odor Coding in the Mosquito. bioRxiv, 2020.11.07.368720. 10.1101/2020.11.07.368720.

65. Schwarzkopf, M., Liu, M.C., Schulte, S.J., Ives, R., Husain, N., Choi, H.M.T., and Pierce, N.A. (2021). Hybridization chain reaction enables a unified approach to multiplexed, quantitative, high-resolution immunohistochemistry and in situ hybridization. Development 148, dev199847. 10.1242/dev.199847.

66. Krüger, A.P., Vieira, J.G.A., Scheunemann, T., Nava, D.E., and Garcia, F.R.M. (2021). Effects of temperature and relative humidity on mating and survival of sterile Drosophila suzukii. Journal of Applied Entomology 145, 789–799. https://doi.org/10.1111/jen.12894.

67. R Core Team (2020). R: A Language and Environment for Statistical Computing.

68. Hothorn, T., Bretz, F., and Westfall, P. (2008). Simultaneous Inference in General Parametric Models. Biometrical Journal 50, 346–363.

69. Benjamini, Y., and Hochberg, Y. (1995). Controlling the False Discovery Rate: A Practical and Powerful Approach to Multiple Testing. Journal of the Royal Statistical Society: Series B (Methodological) 57, 289–300. https://doi.org/10.1111/j.2517-6161.1995.tb02031.x.

70. Rosenthal, R., Cooper, H., and Hedges, L. V (1994). Parametric measures of effect size, in The hand-book of research synthesis. Russell Sage Foundation, 231–244.

