## Supplementary figures for "*Sox21b* underlies the rapid diversification of a novel male genital structure between *Drosophila* species"

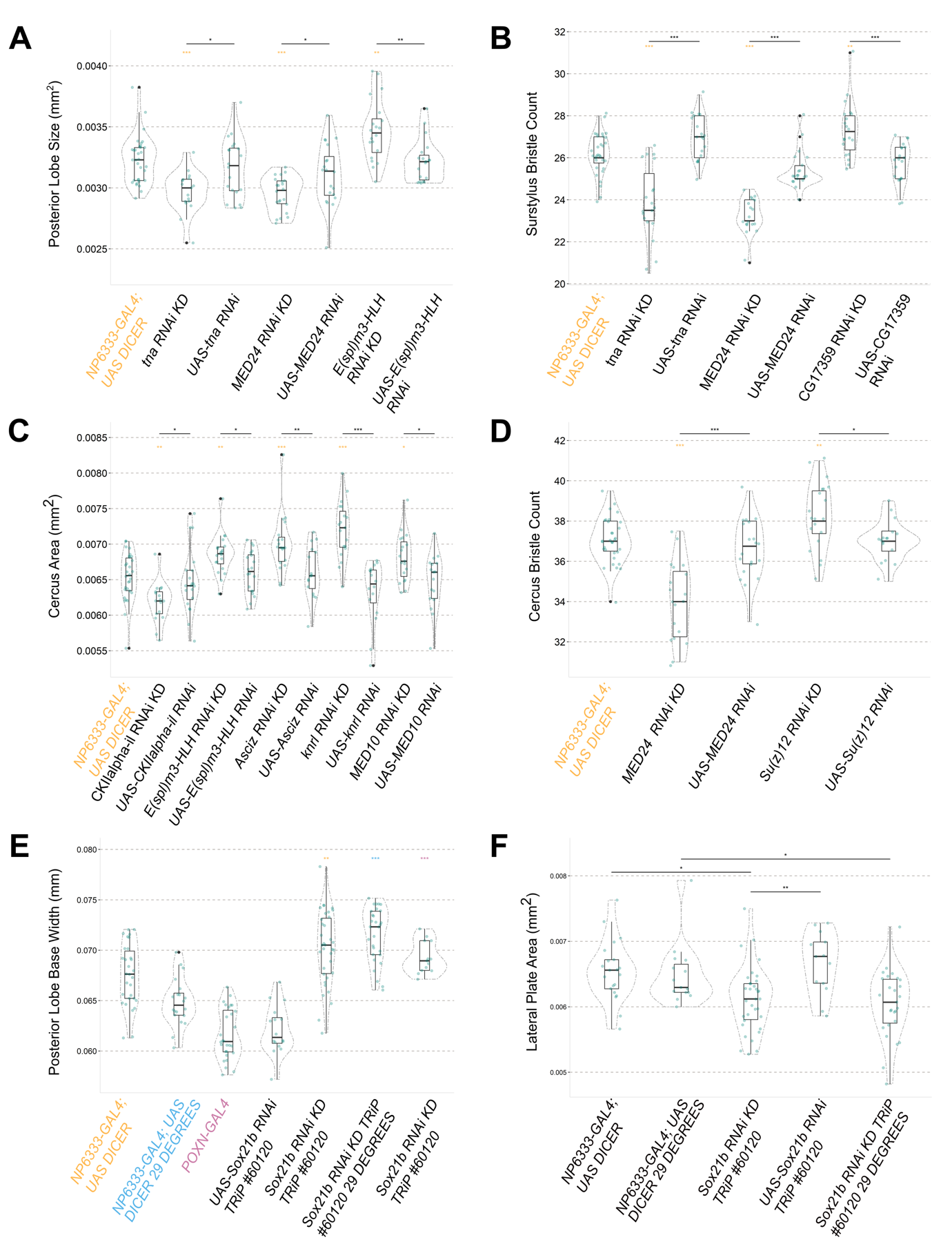

**Figure S1: Genes with a significant effect on periphallic male genitalia structures after knockdown by RNAi in *D. melanogaster.***

(**A-D**) RNAi knockdown in *D. melanogaster* of candidate genes located on chromosome 3, which are differentially expressed between *D. simulans* and *D. mauritiana* developing male genitalia. Knockdowns are classed as significant if statistically different to both parental controls in the same direction. n > 13 for each genotype assayed. Asterisks represent the level of significance (* p<0.05, ** p<0.01, *** p<0.001), is this and all other panels. All RNAi knockdowns were driven by *NP6333-GAL4*. (**E**) RNAi knockdown of *Sox21b* resulted in a significant elongation of the posterior lobe base width. The base width of the posterior lobe was determined at the artificial baseline where it protrudes from the lateral plate. The driver is indicated by the colour of the asterisks above the *Sox21b* RNAi knockdown. For this analysis, only the #60120 TRiP line was used as it revealed a stronger phenotype than #48312 VDRC. (**F**) RNAi knockdown of *Sox21b* showed a decrease in the lateral plate area. The area of the lateral plate was determined as the outline of the plate, adjacent to the posterior lobe, enclosed by an artificial baseline.

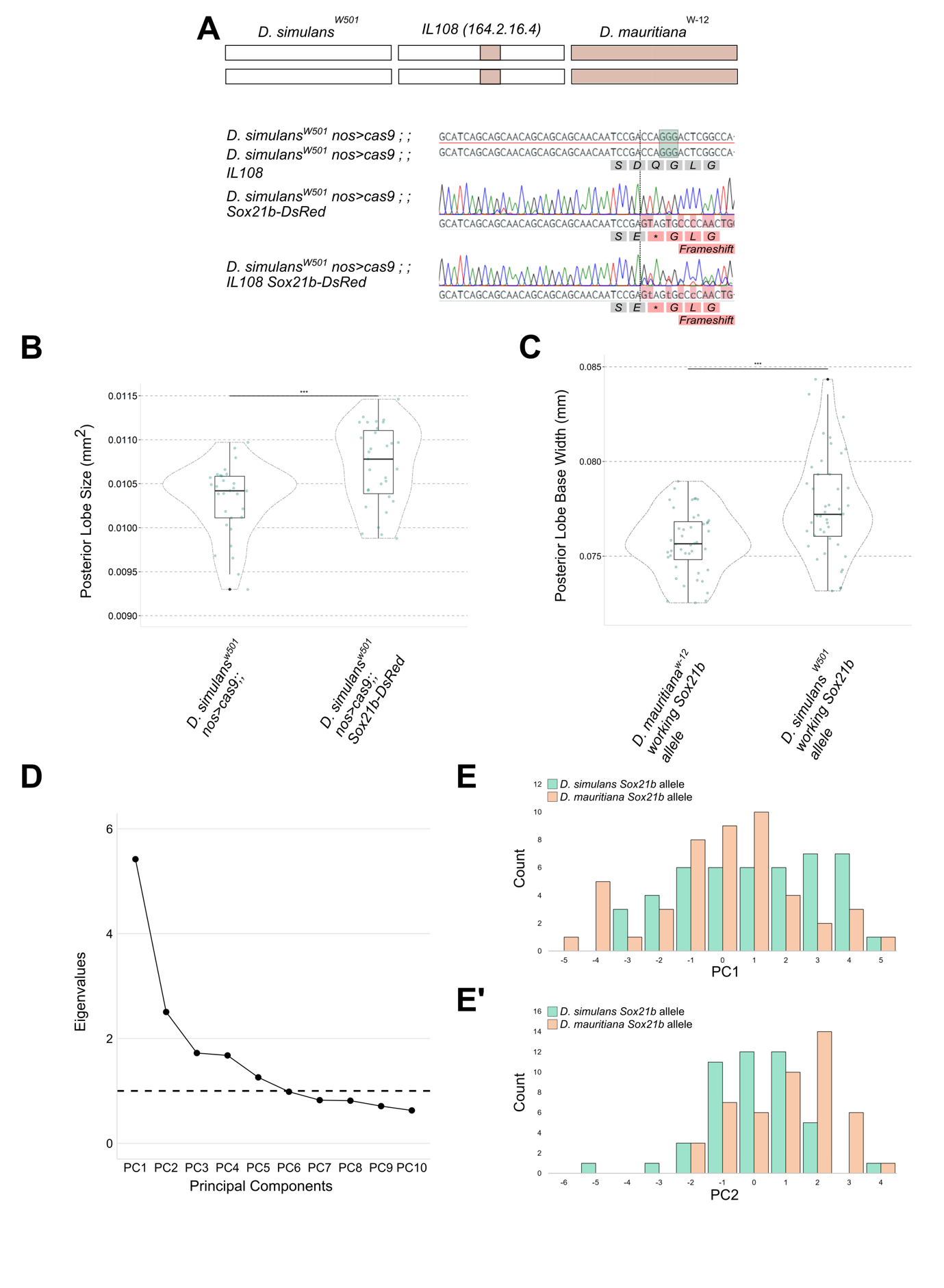

**Figure S2: *Sox21b* posterior lobe shape and size effects in *D. simulans* and the *IL108* introgression line.**

(**A**) Schematic summarising the genetic structure of the *IL108* introgression line on chromosomal arm 3L. Chromatograms illustrating the 3XP3-DsRed insertion compared to the original sequence. The highlighted green box indicates the PAM sequence. The dashed line indicates the CRISPR cut site. Insertion of the pHD-DsRed plasmid induces a frameshift in the open reading frame. (**B**) Homozygous *D. simulans Sox21b-DsRed* flies compared to the non-modified parental line exhibited a significant increase in posterior lobe area. (**C**) Posterior lobe base width is significantly reduced in length when the working *D. mauritiana Sox21b* allele is present compared to the working *D. simulans* *Sox21b* allele (n > 44) (Tables S2 and S4). (**D**) Screenplot representing the Eigenvalue for each principal component (PC). Only the first five PCs had an Eigenvalue >1 and were hence selected for analysis to explore any shape alterations arising from the *Sox21b* allelic origin. Each PC with an Eigenvalue >1 described the following proportion of shape variation: PC1 27.1%, PC2 12.5%, PC3 8.6%, PC4 8.4% and PC5 6.3%. P values indicate the level of significance of PCs when compared between reciprocal hemizygotes using the Wilcoxon Rank Sum Test: PC1 p = 0.02498, PC2 p = 0.000952, PC3 p = 0.6165, PC4 p = 0.739, PC5 p = 0.324. (**E**) Comparison of the shape changes caused by the presence of species-specific working *Sox21b* allele on posterior lobe shape variation summarised by PC1 (p<0.05) and (E’) PC2 (p<0.001).

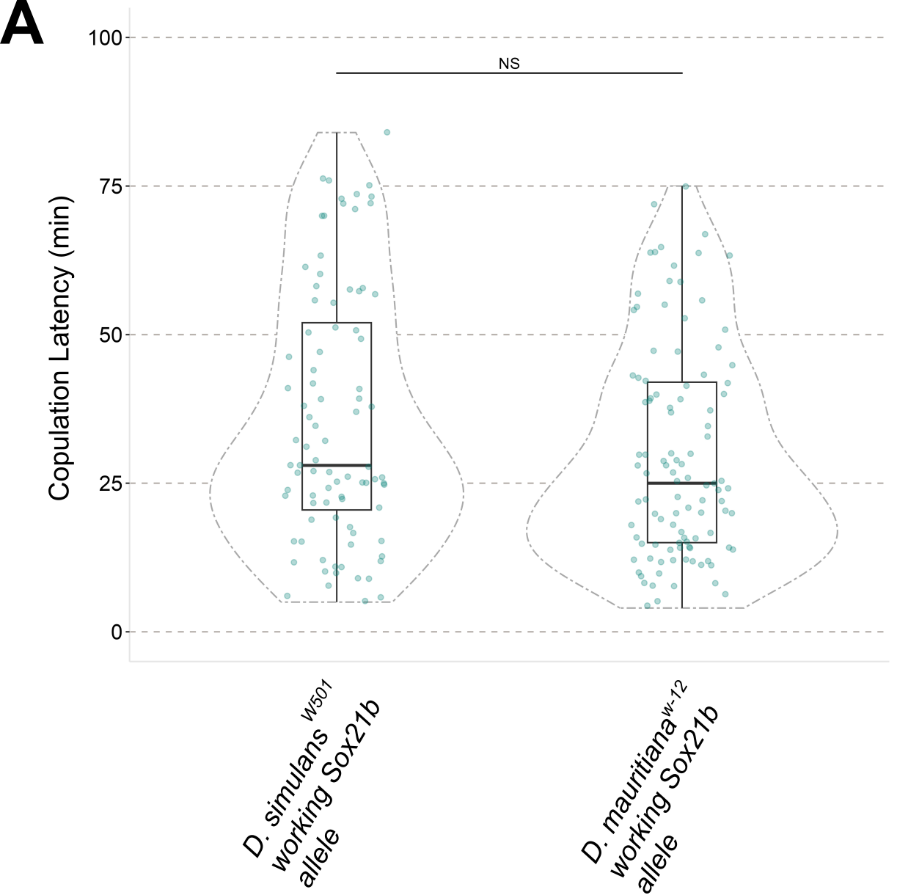

**Figure S3: Comparison of the species-specific *Sox21b* allele effects on copulation latency.**

(**A**) Individual *Sox21b* reciprocal hemizygote males carrying either a working allele from *D. simulans* or *D. mauritiana* were paired with a single *D. simulans* female, but showed no significant difference in copulation latency (n>87).
